# DR-BERT: A Protein Language Model to Annotate Disordered Regions

**DOI:** 10.1101/2023.02.22.529574

**Authors:** Ananthan Nambiar, John Malcolm Forsyth, Simon Liu, Sergei Maslov

## Abstract

Despite their lack of a rigid structure, intrinsically disordered regions in proteins play important roles in cellular functions, including mediating protein-protein interactions. Therefore, it is important to computationally annotate disordered regions of proteins with high accuracy. Most popular tools use evolutionary or biophysical features to make predictions of disordered regions. In this study, we present DR-BERT, a compact protein language model that is first pretrained on a large number of unannotated proteins before being trained to predict disordered regions. Although it does not use any explicit evolutionary or biophysical information, DR-BERT shows a statistically significant improvement when compared to several existing methods on a gold standard dataset. We show that this performance is due to the information learned during pretraining and DR-BERT’s ability to use contextual information. A web application for using DR-BERT is available at https://huggingface.co/spaces/nambiar4/DR-BERT and the code to run the model can be found at https://github.com/maslov-group/DR-BERT.

## Introduction

Over a century ago, the chemist Emil Fischer postulated the lock-and-key model for enzymatic reactions, giving rise to the theory that a protein’s function depends on its unique and rigid three-dimensional structure (***Fischer, 1894***). Within this paradigm, two proteins can interact if they have complementary structures. This idea has contributed to several advances in the understanding of protein function, and it is undeniable that the structure of a protein affects its function. However, studies in the late 1990s and early 2000s recognized that a stable structure is often not necessary for functional function (***Wright and Dyson, 1999; Uversky, 2002***). Segments that lack a rigid structure, also known as intrinsically disordered regions (IDR), have been found in many proteins and shown to actively participate in diverse functions ***Van Der Lee et al***. (***2014***). In fact, these disordered regions are critical for some proteins with central roles in cellular signaling and regulatory networks, allowing them to interact with different proteins (***Wright and Dyson, 1999, 2015***).

Given the functional importance of disordered regions, computational methods for predicting disordered regions have been studied for decades, and over a hundred methods, ranging from biophysical to machine learning-based models, have been developed (***Zhao and Kurgan, 2022***). Recently, predictors that use deep learning have gained traction (***Zhao and Kurgan, 2022***). This was particularly evident in the Critical Assessment of protein Intrinsic Disorder (CAID) competitions, where deep learning-based models consistently delivered the best performance (***Necci et al., 2021***). Many existing deep learning methods to predict disordered regions utilize recurrent neural networks and convolutional neural networks, sometimes paired with an attention mechanism (***Hanson et al., 2019; Tang et al., 2022, 2020***). This success of deep learning-based methods to predict disordered regions in proteins can be attributed to both the complex and non-linear nature of sequence-structure maps as well as the steady increase of data availability (***Piovesan et al., 2016***).

Protein language modeling has been a particularly fast-growing area of deep learning research for computational biology. Inspired by natural language processing, the core idea of protein language modeling is that the amino acids (or sometimes small groups of amino acids) that make up a protein are analogous to the words that make up a sentence (***Rives et al., 2021; Nambiar et al., 2020***). Like their natural language counterparts, protein language models leverage the large number of unannotated amino acid sequence data to pretrain deep learning models before specializing them on much smaller amounts of annotated data. Usually, this pretraining step consists of training the model either to predict the context surrounding a particular residue (***Asgari and Mofrad, 2015***) or to predict the identity of a hidden residue, given its surrounding context, i.e. the set of its nearby amino-acid residues (***Rives et al., 2021***). These models have then been successfully used to perform various downstream tasks including protein family labeling (***Asgari and Mofrad***, ***2015; Nambiar et al., 2020***), prediction of protein interactions (***Nambiar et al., 2020***) and subcellular localization (***Stärk et al., 2021***), and the inference of evolutionary trajectories and phylogenetic relationships of proteins (***Hie et al., 2022; Lupo et al., 2022***). While most protein language models tend to be large and GPU intensive, there have been studies proposing small and computationally inexpensive protein language models (***Nambiar et al., 2020***).

In this paper, we present Disordered Region prediction using Bidirectional Encoder Representations from Transformers (DR-BERT), a small protein language model that is first pretrained on a large corpus of amino acid sequences and then finetuned to predict disordered regions in proteins. We validate our model on both CAID 1 and CAID 2 evaluation data and benchmark it against some of the best performing models. We then investigate the impact of pretraining on the performance of DR-BERT. Finally, we dive into one particular biological case study involving RPB6, a subunit of RNA polymerase, to illustrate how DR-BERT arrives at its predictions and learns to use contextual information from the amino acid sequence.

## Results

While many models for disordered region prediction depend on knowledge of biophysical properties of amino acids used as inputs, previous work has shown that pretraining a protein language model may allow it to learn these biophysical and functional properties in a self-supervised manner (***Rives et al., 2021; Nambiar et al., 2020***). Therefore, we chose to build our DR-BERT model using only the amino acid sequence of a protein as the input. This model is first pretrained on the masked language modeling task as shown in Figure 1 before it is finetuned to predict intrinsically disordered regions.

**Figure 1.**
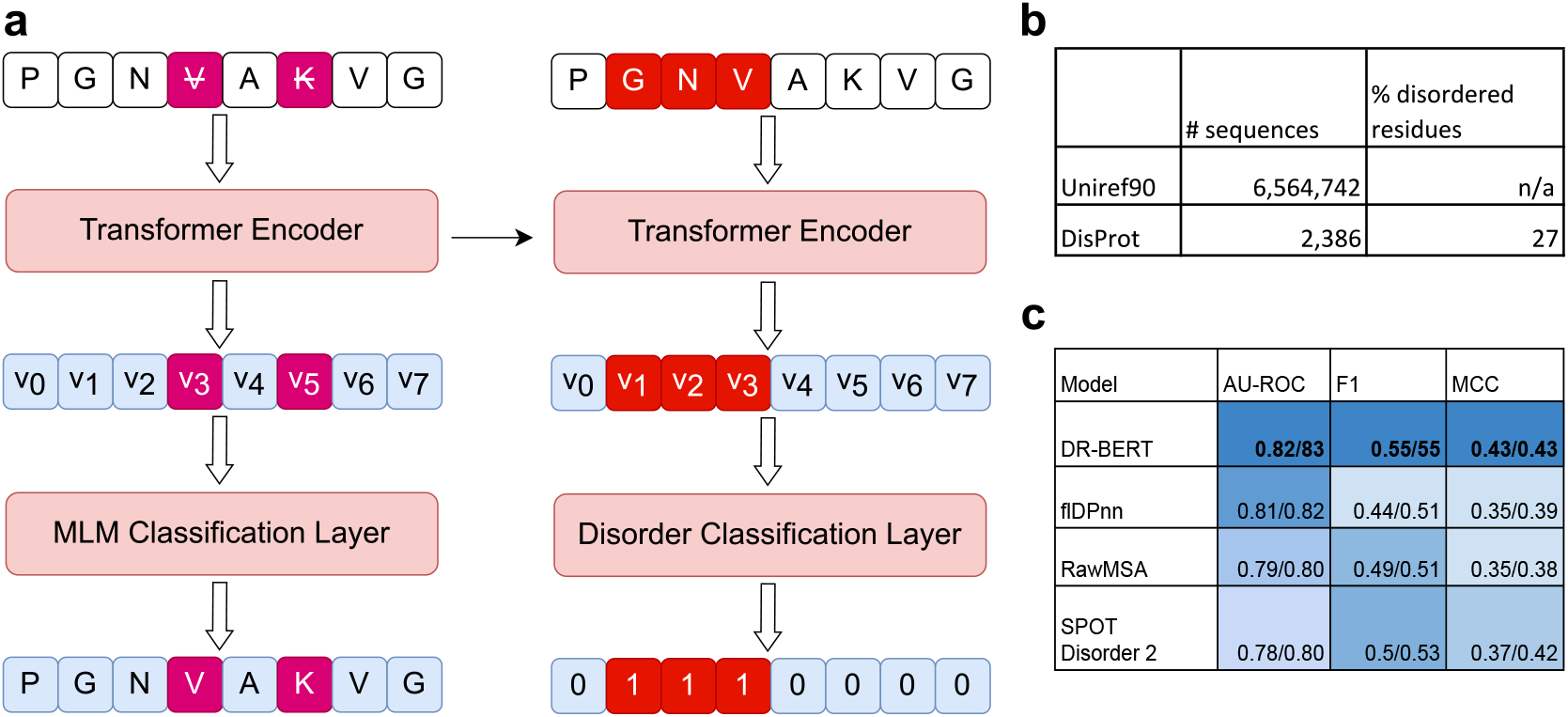
The DR-BERT model is pretrained on the masked language modelling task and finetuned on predicting disordered regions in proteins. **(a)** A schematic of the DR-BERT model and the pretraining and finetuning procedures. **(b)** The statistics of data used in this study and **(c)** The CAID 1 and CAID 2 results of DR-BERT compared to some of the best-performing models from the CAID competitions (***Necci et al., 2021***). Cells are colored based on the performance of each model for a particular metric for CAID 1.

The model itself is a neural network with a Transformer encoder block composed of six stacked Transformer encoder layers (see Methods for details). The purpose of the encoder block is to create contextual latent representations of each residue. That is, each residue is represented by a vector that captures the context of the rest of the sequence. By stacking multiple transformer encoder layers within the encoder block, the final latent representations can capture more complex higher-level information and relationships from the amino acid sequence. These vectors are then passed to a final linear layer that constructs a task-specific output.

In the pretraining task of masked language modeling, the neural network is asked to predict the identities of amino acids that have been masked in the input. In this study, we pretrained our model on 6,564,742 proteins randomly sampled from the UniRef90 dataset (***Suzek et al., 2014***). Next, we finetuned DR-BERT by tasking it to classify residues in proteins as disordered or ordered using annotated data from the DisProt database. The performance of DR-BERT on this finetuning task is shown in Figure 1c alongside previous state-of-the-art methods.

### Benchmarking DR-BERT’s performance

When finetuning DR-BERT on the disordered region classification task, we split the DisProt data into train/validation/test sets with the aim of enabling a systematic and unbiased comparison against existing methods. In particular, proteins from the Critical Assessment of Protein Intrinsic Disorder Prediction (CAID) competitions were reserved as test data and were not available to the model during training. In addition, any proteins that shared more than 25% similarity to proteins from the test set was excluded from the train set. We ran out our benchmarking on both CAID 1 and CAID 2. This left us with 1,408 examples in the train set, 156 sequences in validation and 652 in the test set for CAID 1 and 2013 examples in the train set, 216 sequences in validation and 348 in the test set for CAID 2. By doing so, we were able to reproduce the results of some of the top-performing models from CAID (***Necci et al., 2021***). In particular, we benchmarked DR-BERT against flDPnn (***Hu et al., 2021***), RawMSA (***Mirabello and Wallner, 2019***), SPOT-Disorder2 (***Hanson et al., 2019***), DisoMine (***Orlando et al., 2022***), Espritz-D (***Walsh et al., 2012***), AUCpreD (***Wang et al., 2016***), IUPred2A/3 (***Mészáros et al., 2018***) and Predisorder (***Deng et al., 2009***).

Of these methods, all but IUPred2A/3 are deep learning-based models based on feed-forward, recurrent, and convolutional neural network architectures. flDPnn is a feed-forward neural network that uses evolutionary and structural information in addition to disordered region predictions from simpler models; RawMSA uses convolutional neural networks (CNNs) on evolutionary information (in the form of MSAs); SPOT-Disorder2 uses a combination of CNNs and recurrent neural networks (RNNs) on input with evolutionary information; DisoMine uses RNNs on structural information; Espritz uses RNNs on evolutionary information; AUCpreD uses CNNs on sequence information (with optional evolutionary information); and Predisorder uses RNNs with structural, biophysical and evolutionary information. A notable pattern here is that most of these methods use pre-computed features. First, the performance of the model is reliant on its upstream dependencies. For example, if a model uses MSAs as input, one would expect its performance to deteriorate for proteins that do not have many known homologs. In addition, the presence of multiple third-party techniques in a prediction pipeline makes it more difficult to optimize computational efficiency. In contrast, DR-BERT is a fully self-contained model that does not rely on any additional information besides the amino acid sequence of a protein.

Despite not requiring any additional information, the Receiver Operating Characteristic (ROC) curves (Figure 2) on both the CAID 1 and CAID 2 test sets demonstrate that DR-BERT outperforms all of the other methods in predicting disordered regions. For CAID 1, DR-BERT ranks first in terms of area under the ROC curves (AU-ROC) with a value of 0.82. The scores then incrementally decrease with flDPnn, RawMSA and SPOT-Disorder2. For CAID 2, DR-BERT is again the highest ranking method followed by flDPnn and a three way tie between rawMSA, SPOT-Disorder2 and Disomine. The ROC curves also show that DR-BERT offers particularly evident improvements in the lower range of false positive rates. However, as the disordered region dataset is imbalanced with more ordered residues than disordered ones, the ROC curves may show an overly optimistic view of the classifiers (***Davis and Goadrich, 2006***). Therefore, we also calculate F1 and Matthews correlation coefficients (MCCs) for each model. These scores, along with the AU-ROC scores are shown on Figure 3a for CAID 1 and Figure 3b for CAID 2. Again, DR-BERT scores the highest on both metrics with an F1 of 0.55 and MCC of 0.43 for CAID 1 and an F1 of 0.56 and MCC of 0.43 for CAID 2. The precision-recall plots on Supplementary Figure 1 also shows that DR-BERT performs better than the other methods in balancing the trade-off between precision and recall.

**Figure 2.**
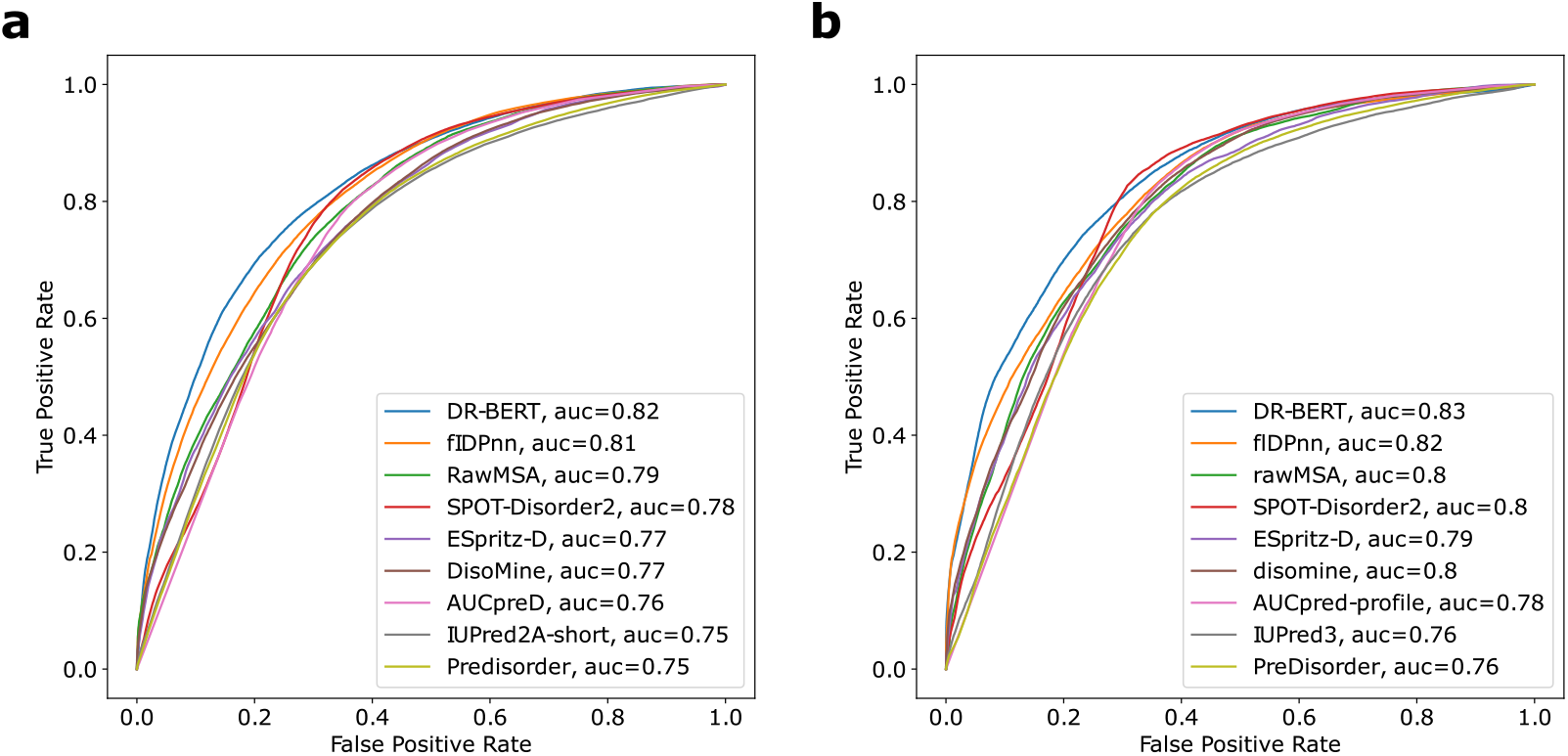
The ROC curves of DR-BERT and other models on test sets from (a) CAID 1 and **(b)** CAID 2. The legends display the area under the curve (AUC) for each model. The models are ordered based on the AUC in CAID 1.

**Figure 3.**
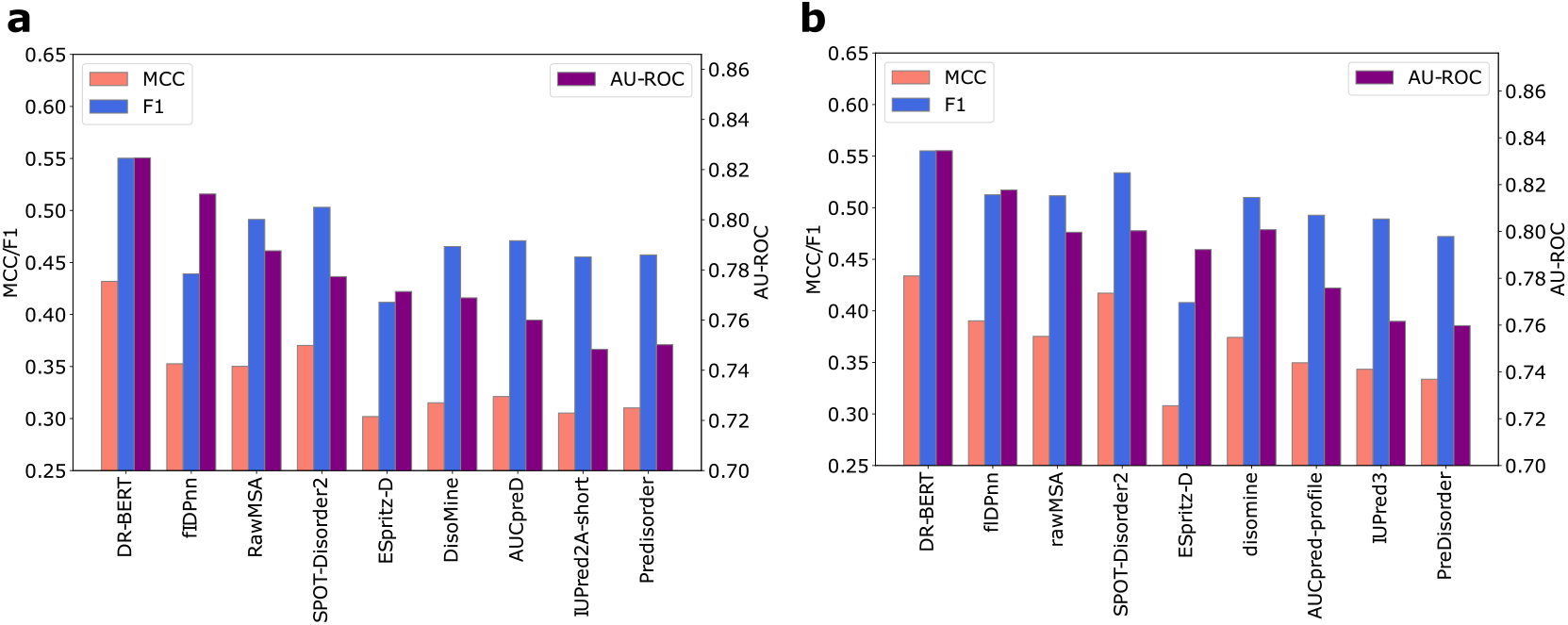
Comparing the results of DR-BERT with other top-performing methods on the CAID datasets. **(a)** The MCC, F1, and AU-ROC scores of DR-BERT and the top performing methods from CAID 1, evaluated on the test split. **(b)** The MCC, F1, and AU-ROC scores for corresponding methods evaluated on the CAID 2 test data.

To determine the statistical significance of DR-BERT’s improvement over the existing methods, we performed a resampling analysis similar to that of ***Hu et al***. (***2021***) and ***Necci et al***. (***2021***). Specifically, we resampled 25% of the test set 20 times. For each resample, we calculated the AU-ROC, F1, and MCC scores for DR-BERT and the other methods. Finally, we performed Wilcoxon tests comparing the scores obtained by DR-BERT to those of each of the other methods, with the alternative hypothesis being that DR-BERT’s score is greater than the other method. The *p*-values from these hypothesis tests, shown in Supplementary Figure 2, confirm that DR-BERT performs significantly better than existing methods across the different scoring metrics for both CAID 1 and CAID 2.

This supremacy of DR-BERT over methods that use evolutionary and structural features suggest that these features can be successfully learned by the model either during pretraining or finetuning. In fact, it has been previously shown that pretrained protein language models are able to extract structural information from amino acid sequences (***Bhattacharya et al., 2020; Singh et al., 2022***). However, these results alone do not elucidate the contribution of pretraining to the success of DR-BERT.

### Pretraining and Finetuning

To better understand the role that pretraining plays in extracting the information relevant to disordered region prediction, we interrogated DR-BERT models at two stages: (a) after only pretraining and (b) after pretraining and finetuning. At both of these stages, we extracted the embeddings from the Encoder Block for each residue in the test set. Using t-SNE, we projected these embeddings down to two dimensions (***Van der Maaten and Hinton, 2008***). Then, we calculated kernel density estimates (KDEs) separately for ordered and disordered residues. These KDEs are shown in Figure 4a for the pretrained model and in Figure 4b for the model that was pretrained and fine-tuned. The plot for the pretrained model shows about 20 different clusters of ordered residues and 15 distinct clusters of disordered residues. Upon further investigation, we see that each cluster corresponds to an individual amino acid. There are a few exceptions to this for disordered residues. For instance, there is no clear disordered cluster for the amino acid tryptophan (W). This is because tryptophan is one of the most order-promoting amino acids and is rarely encountered inside intrinsically disordered regions (***Campen et al., 2008***). However, the clear overall pattern in Figure 4a is that most ordered clusters are accompanied by an adjacent disordered cluster for the same amino acid. This is in contrast to the null model where the disorder/order residue labels are shuffled, shown on Supplementary Figure 5. On the other hand, the plot for the finetuned embeddings depicts a different story. While the embeddings are not clustered by amino acid, the disordered residues are all clustered together and are well-separated from the ordered residues. The difference between the pretrained and finetuned embeddings highlights that pretraining a protein language model is sufficient to extract some information regarding disordered regions in proteins. Finetuning the model allows it to then hone in on the differences between disordered and ordered residues to more efficiently separate them. This result gives credence to an observation we made in ***Nambiar et al. (2020***) where we noted that pretraining a protein language model allows it to learn general but biologically relevant information from amino acid sequences, whereas finetuning gives the model more information about one characteristic but at the expense of generality.

**Figure 4.**
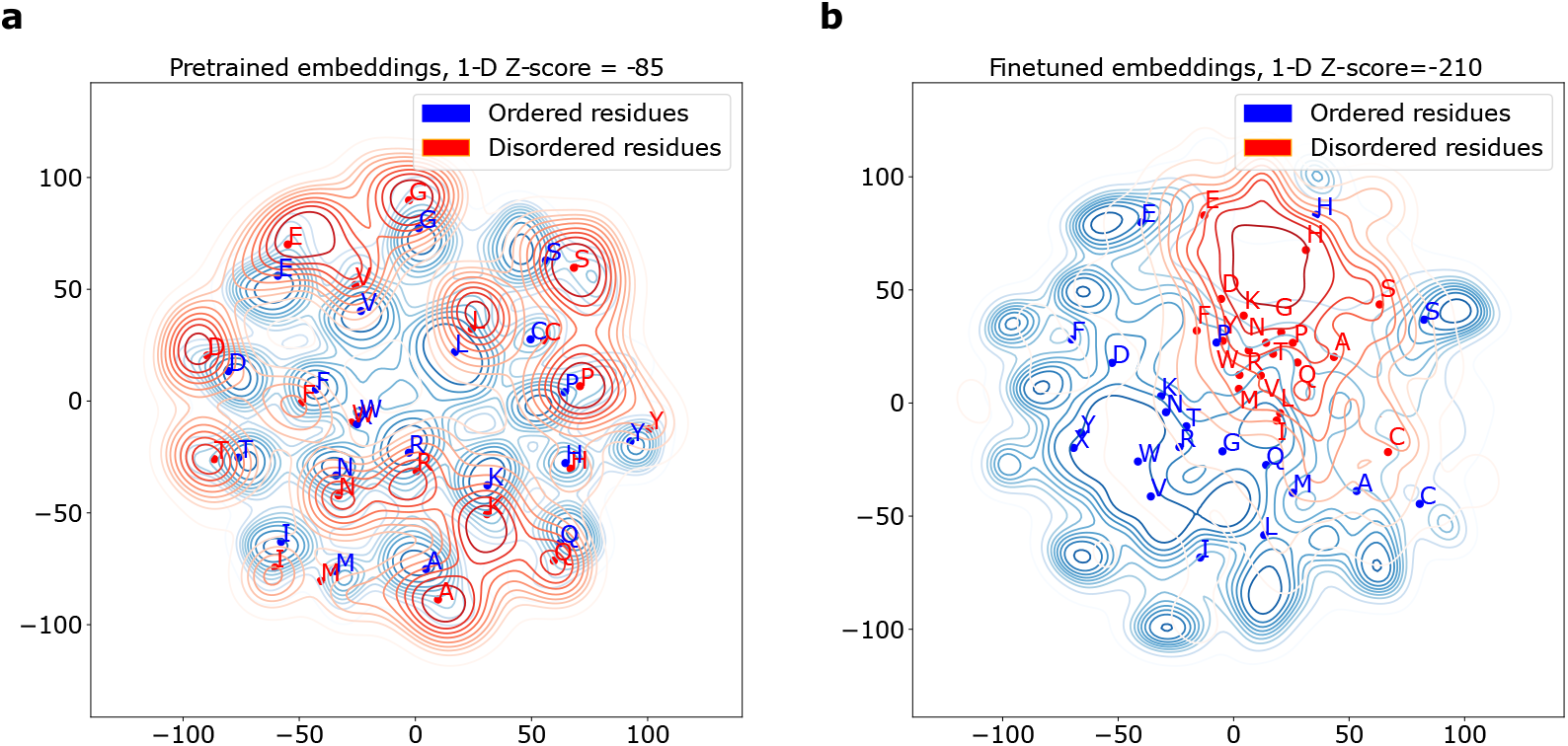
Plotting the embeddings of ordered and disordered residues. **(a)** A t-SNE projection of the pretrained embeddings of residues in the CAID 1 test set. The plot shows the kernel density estimates of ordered residues in blue and disordered residues in red. The labeled points indicate the mean position of each amino acid. This plot should be compared to the null model shown in Supplementary Figure 5. **(b)** A similar plot but with embeddings from a model finetuned to predict disordered regions. For both plots the two sample Z-test is performed after reducing the dimensionality of the embedding to 1-D.

In addition, we wanted to quantify the advantage of pretraining for predicting disordered regions. To do this, we trained a model with an identical architecture to DR-BERT to predict disordered regions without any pretraining. The results of this non-pretrained model evaluated on CAID 1, shown in Figure 5, show that pretraining DR-BERT gives it a considerable advantage. In fact, in the absence of pretraining, our model lags behind the models from the CAID competition, shown in Figure 3 and Supplementary Figure 4. This showcases the advantage of pretraining for Transformer neural networks, especially for low data regimes.

**Figure 5.**
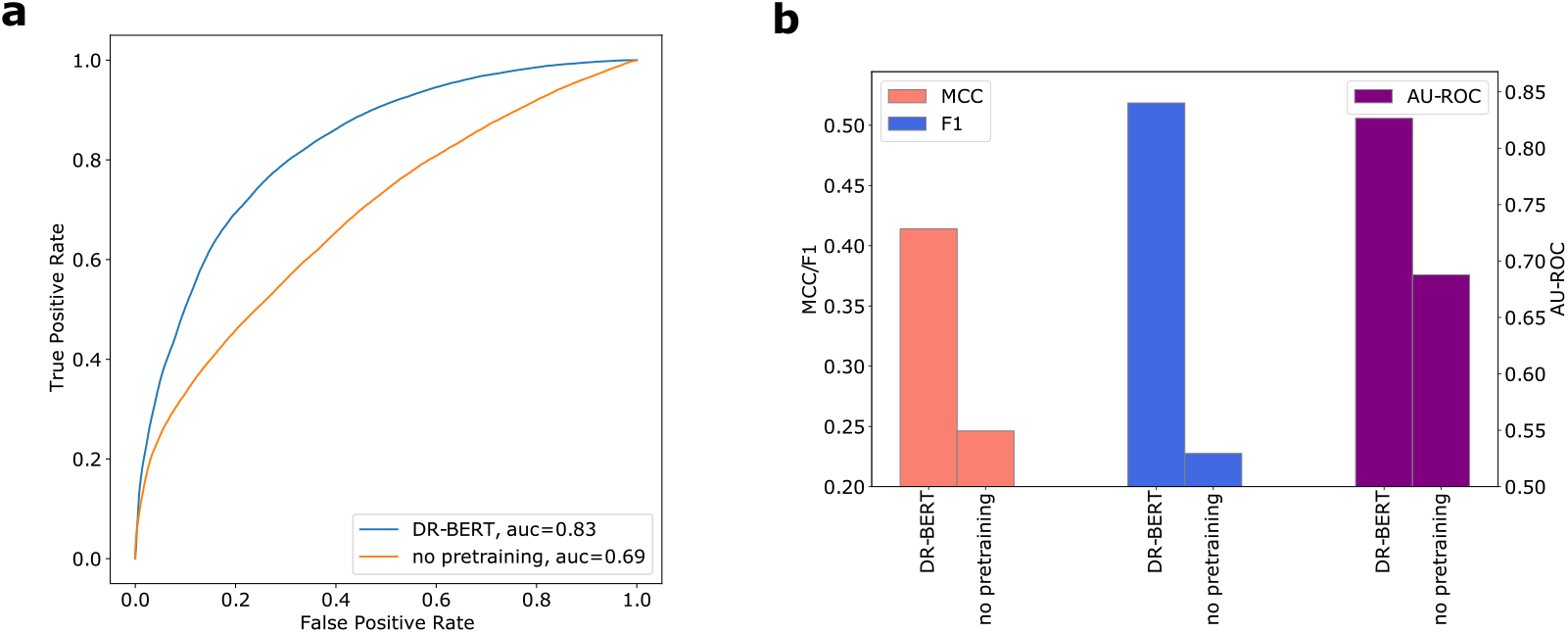
Comparison of the results of DR-BERT with its version without pretraining. **(a)** The ROC plots of DR-BERT and the non-pretrained model, evaluated on the CAID 1 test set. The area under each curve (AUC) is presented in the legend. **(b)** The MCC, F1 and AU-ROC scores of DR-BERT and the version without pretraining.

### A case study: the disordered region in RPB6 protein

Evaluating DR-BERT on a large annotated dataset gave us confidence in DR-BERT’s ability to make accurate predictions regarding disordered regions. However, it is also useful to illustrate a potential use-case by focusing on the predictions of the model for an intrinsically disordered region within a single protein. Doing so gives us the opportunity to gain insight into how the context of a particular sequence is used by the attention heads of the model (see Methods) to make predictions for different residues in the same protein. We decided to illustrate this using RPB6 protein as an example. RPB6 is a subunit of an RNA polymerase in fission yeast. It is known to bind to the general transcription factor, TFIIS (***Ishiguro et al., 2000***). This example allows us to test our disordered region prediction for a protein that is known to perform an important function. Figure 6b shows that DR-BERT predicts with high confidence that the N-terminal tail of RPB6 is in fact disordered. Indeed, NMR spectroscopy shows that not only does RPB6 have a flexible N-terminal tail, this tail is also used to bind to the p62 subunit of the TFIIH transcription factor(***Okuda et al., 2021***). Figure 6a shows DR-BERT’s predictions overlaid on the NMR-determined structure of the complex between RPB6 and the TFIIH p62 PH domain (PDB: 7DTI). To analyze how DR-BERT uses sequence context to make its predictions, we extracted the self-attention heads for each of the six layers in DR-BERT’s encoder as it processed the 130 amino acid long RPB6 sequence. Each attention map is represented as a 130 × 130 matrix *M* where *M*_*ij*_ gives a numerical score as to how much that particular attention head focuses on amino acid *j* when determining the context relevant for amino acid *i*. A sample of these attention maps for each layer in DR-BERT is displayed in Figure 6c (a complete table is shown on Supplementary Figure 6). We observed that the features learned by the attention maps of the initial layer do not have any clear high-level patterns. However, attention maps from layers two to four display some distinct patterns. For example, in layer 4, the attention map reveals that the relevant context for each residue includes a large window of surrounding residues in addition to several smaller windows at intervals on either side of the residue in question. By layer 5, at least one attention map divides the residues into two distinct groups: one group consists of residues 1 to 50, and the other group comprises residues 50 to 130. Each group only considers residues within its respective group as relevant context, while ignoring residues in the other group. Comparing this attention map to the disorder scores by sequence position, we can see that the division between the two groups occurs at the transition between the ordered and disordered regions of the protein. This observation is similar to demonstrations in computer vision, where deep neural networks learn features hierarchically, with the initial layers detecting simpler, disjoint features, and layers towards the end of the neural network detecting high-level features directly related to the model’s training task.

**Figure 6.**
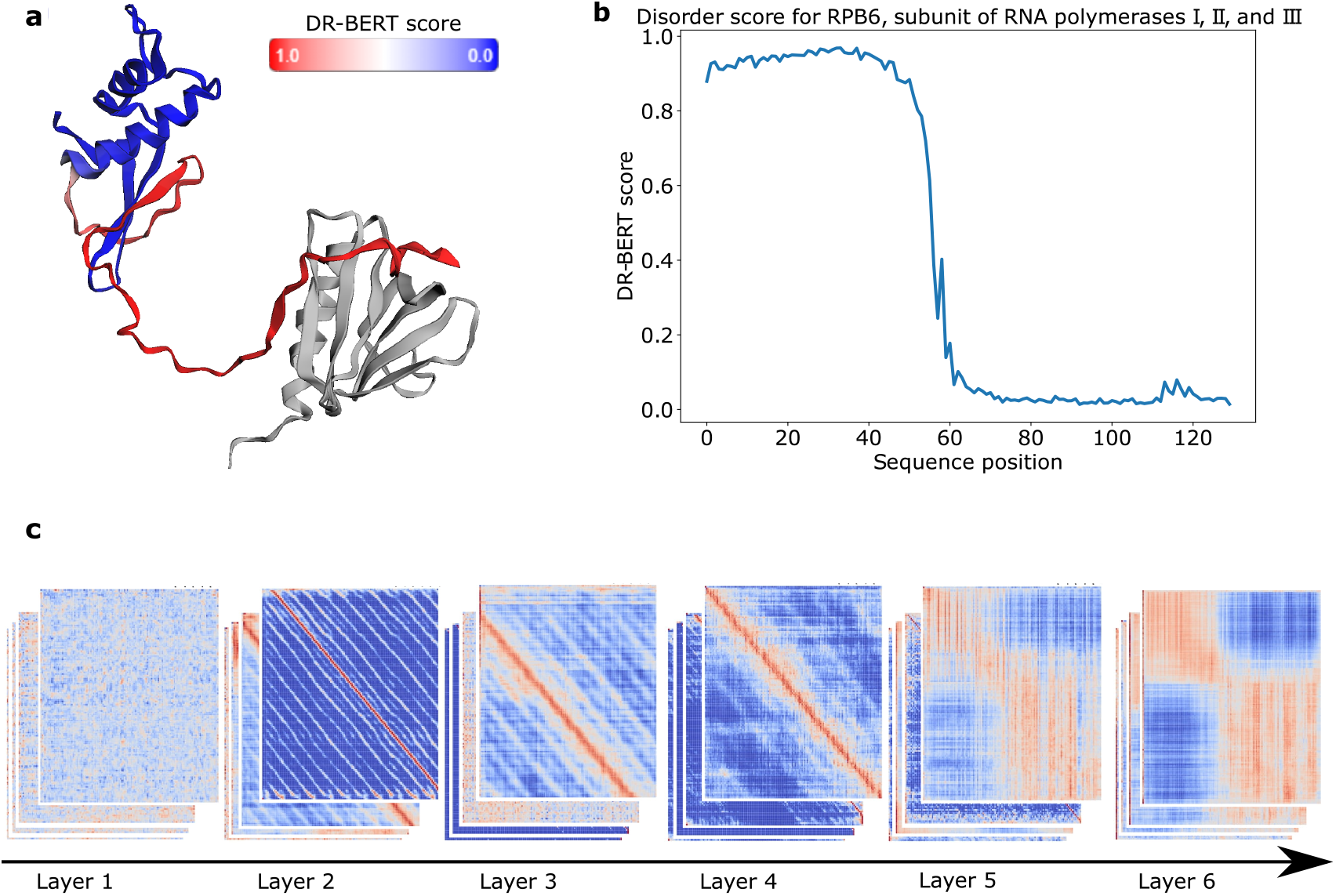
Application of DR-BERT to RPB6, a subunit of RNA polymerase. **(a)** The three-dimensional structure of RPB6 as it binds to the TFIIH p62 PH domain (PDB:7DTI). The protein is colored by the DR-BERT score, which represents the probability that a given residue is disordered. **(b)** A plot of the DR-BERT scores for RPB6 shown for each position along the amino acid sequence. **(c)** A sample of DR-BERT’s self-attention maps for each of the 6 layers in the model as it processes the RPB6 sequence. The attention maps have been log transformed, and the red cells indicate higher attention values while the blue cells indicate lower attention values.

## Discussion

In this study we introduce DR-BERT, a protein language model for predicting disordered regions in proteins. DR-BERT is first pretrained on the masked language modeling task before it is finetuned to predict disordered regions.

This finetuned model is benchmarked using the CAID evaluation data and significantly surpasses the other models. This improvement over models that use biophysical and structural information supports the hypothesis that pretraining protein language models enables them to learn biologically relevant information in a self-supervised manner without any provided annotations.

This hypothesis is further validated as we show that the embeddings of the pretrained model are able to differentiate between disordered and ordered residues without access to any annotations during training. Furthermore, we showed that a model with an identical architecture as DR-BERT suffers a large loss in performance when the pretraining step is skipped.

Finally, we took a closer look at how DR-BERT makes predictions for RPB6. Through this exercise we saw that DR-BERT extracts patterns hierarchically, with higher level features extracted by attention heads in deeper layers of the neural network. This is similar to behavior that had been observed in computer vision.

To verify that DR-BERT was not overfitting on the training data, we excluded from the training set proteins that were clustered with proteins in the test set with 25% sequence similarity. The clustering in this process was performed using CD-HIT (***Fu et al., 2012***).

Given the high performance of DR-BERT on the disordered region prediction task, we also investigated its ability to perform related tasks from CAID 2. This included evaluating DR-BERT on a disordered region dataset where X-ray attotations were removed (disorder-noX) and a dataset where PDB residues were incorporated (disorder-PDB). The results shown on Supplementary Figure 3 show that DR-BERT performs well on the disorder-noX test set, placing first on the AU-ROC and area under precision-recall plot metrics and second to SPOT-Disorder2 on F1 score and MCC. However, DR-BERT only shows middling performance on the disorder-PDB set. Given that DR-BERT was trained on the vanilla disordered region dataset from Disprot, it is not surprising that DR-BERT’s performance dropped on some of these variants. In addition to variants of disordered region annotations, we also evaluated DR-BERT on predicting protein-binding regions. Protein-binding disordered regions are regions in disordered proteins that bind to structured partners and potentially allow the disordered protein to bind to multiple partners (***Mészáros et al., 2009***). DR-BERT achieved an AU-ROC of 0.75, F1 score of 0.45 and MCC of 0.32 beating other protein-binding predictors from CAID 2.

The success of DR-BERT, in addition to the insight into how DR-BERT makes predictions, leads us to believe that protein language models could play an important role in the next generation of neural networks for predicting disordered regions. In fact, after completing our study, we found that a similar model to DR-BERT was presented in a recent preprint by ***Redl et al***. (***2023***). However, there are significant differences in our studies, including our investigation of the effect of pretraining on the success of the protein language model and the insight into the features extracted by the attentional layers. In addition, DR-BERT is significantly smaller than the model proposed by ***Redl et al***. (***2023***) (with 15x fewer parameters), which may make DR-BERT more accessible to users without access to high-performance GPUs. An alternative approach to the one shown in our study would be to extract embeddings from a pretrained model and pass them to a downstream classifier without finetuning the embeddings. This approach, which is used by the SETH model, makes it more efficient to train models on downstream tasks using embeddings from a large pretrained language model (***Ilzhöfer et al., 2022***). However, as shown in Supplementary Figure 7, DR-BERT is able to surpass the performance of a larger language model where the embeddings are not finetuned. Moreover, each prediction would still involve a forward pass on the large model, which would be slower than using a small protein language model like DR-BERT.

In order to maximize the accessibility of our model, we have made a web-app (accessible at https://huggingface.co/spaces/nambiar4/DR-BERT) where anyone can use DR-BERT to make disordered region predictions. Due to the small size of our model, we are currently able to run our server using only 2 CPU cores and 16GB of RAM. In addition to the web-app, users who want more control over their predictions can run the pretraining, finetuning, and prediction scripts we have made available at https://github.com/maslov-group/DR-BERT.

While we have shown that a protein language model with no additional information is sufficient to make accurate predictions of disordered regions in proteins, a direction worth exploring in the future is whether combining the information learned by protein language models with biophysical properties and the outputs of other models might further improve performance. One additional input that could be particularly interesting is the per-residue confidence score provided by AlphaFold when making sequence-to-structure predictions since it has been shown that disordered regions are often assigned a low confidence score by AlphaFold (***Alderson et al., 2022***).

## Methods and Materials

### Data Processing

#### Pretraining Data

From UniRef90, we sampled 6,564,742 proteins at random for the training dataset and 250,000 proteins for the validation dataset. Per the construction of UniRef data, the validation set is designed to contain no examples with above 90% sequence similarity to any of the examples in the training set (***Suzek et al., 2014***). We used Uniref90 because we hope that down the road, we can use DR-BERT to predict the effects of mutations on disordered regions. It has been observed in ***Meier et al***. (***2021***) that the redundancy provided by Uniref90 allows for better variant effect prediction.

#### Disordered Region Data

2,419 sequences were taken from DisProt Version 9.2, June 2022, with replicate proteins removed to create the 2,386 example dataset, and their disordered regions were recorded into boolean arrays as the ground truth labels (***Quaglia et al., 2021***). To construct the train/validation/test splits for CAID 1, the Disprot and CAID 1 datasets were combined and clustered for 25% similarity using the CD-HIT algorithm. Then, only clusters without any CAID 1 proteins were used for training and validation. This resulted in 1,569 proteins in the train set and 156 sequences in the validation set. The 652 proteins from the CAID 1 competition dataset were used as the test set. To construct the train/validation/test splits for the CAID 2 experiment, we similarly combined the CAID 2, CAID 1 and Disprot datasets, and clustered for 25% similarity. Then, only clusters without any CAID 2 proteins were used for training and validation. The sizes for the train, validation, and test sets were 2013, 216, and 348 respectively. In addition to the standard disorder dataset from CAID 2, we also tested on a test set were X-ray annotations were removed (disorder-noX), a test set where PDB residues were incorporated (disorder-PDB) and a test set for protein binding. These results are included in Supplementary Figures.

### Model Architecture

#### Embedding Layers

DR-BERT uses the Bidirectional Encoder Representations from Transformers (BERT) coupled with token classification heads trained on disordered region labels (***Devlin et al., 2018***). Based on the Robustly Optimized BERT pretraining Approach (RoBERTa), the model consists of an embedding layer connected with a Encoder Block with 6 encoder layers. The embedding layer consists of two main component layers: a word embedding layer and a positional embedding layer. The word embedding layer takes the tokenized sequence of amino acids and maps each token to a 768 dimensional vector. In contrast, the positional embedding layer captures the spatial information of the tokens to preserve the notion of context within the sequence (***Vaswani et al., 2017***).

#### Encoding Layers

After a dropout layer is applied to decrease the potential for overfitting (***Srivastava et al., 2014***), the embedding, consisting of a 768 dimensional vector for each amino acid token, is used by the Transformer encoder layers. The RoBERTa transformer layer consists of a self-attention layer and a feed-forward network layer. The self-attention mechanism described in (***Vaswani et al., 2017***), captures the relationship between different tokens in a sequence. Each attention layer consists of 12 heads, which can each capture different contextual information in parallel. The final output from the encoder layers is 1026 vectors, each of length 768, where the first corresponds to a standard summary [CLS] token and the last corresponds to a [SEP] separator token. Many of the hyperparameters used in this paper, including the hidden size of 768 and 12 attention heads, are based on our previous work in ***Nambiar et al***. (***2020***).

### Pretraining

Pretraining of DR-BERT used masked language modeling (MLM): in each example, the model is tasked with identifying some hidden tokens. Following RoBERTa, the masks are set independently during epochs, and 15% of tokens are replaced with a [MASK] token for each example, with crossentropy loss being applied for every batch of proteins (***Liu et al., 2019***). Pretraining lasted for approximately 11 epochs, allowing the model to see 70 million examples. The batch size was set to 10 examples per device, and the model was trained on 2 NVIDIA V100s.

### Disordered Region Prediction

To finetune DR-BERT, we applied a token classification training method. A classification layer is trained and applied to each positional embedding output Then, a softmax function is applied to transform the embedding into probability space, taking the rounded result as the predicted label. Then, cross-entropy loss is applied between the predicted labels and the ground truths. The classification training lasted 10 epochs, with the best-performing checkpoint on the validation dataset chosen as the final model. The learning rate was empirically chosen to be 2*e*^−6^ (against 2*e*^−5^ and 2*e*^−7^) using the cosine scheduler with hard restarts, as opposed to a linear scheduler. To compare against similar models, DR-BERT was tested on the CAID dataset, which we ensured to be disjoint from both the training and evaluation datasets. We also tested on the CAID 2 dataset, which was again ensured to be disjoint from the training and evaluation datasets.

### Evaluation Metrics

The primary evaluation metrics used for DR-BERT were Area Under the Receiver Operating Characteristic Curve (AU-ROC), F1 score and the Matthews Correlation Coefficient (MCC). The receiver operating curve is given by observing the change in the true positive to false positive ratio as the probability decision threshold is varied. Therefore, ROC-AUC for a perfect classifier would be 1.0 and a random classifier would have an area of 0.5. F1 scores are computed as a flattened vector of all predicted disorder binary labels against their ground truth, and is given by

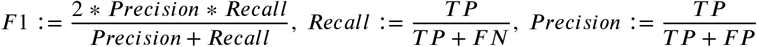

The MCC score offers a metric that is stable in imbalanced datasets (***Chicco and Jurman, 2020***). Because of the MCC formula’s false-positive symmetry, the MCC metric is invariant on which class is considered to be negative or positive. As the DisProt dataset has approximately 3 times as many ordered labels as disordered, the MCC metric is an appropriate metric to characterize the model’s performance. MCC is defined as:

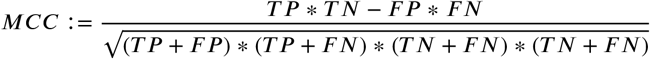

For a fair comparison between methods, when these evaluation metrics were run, only test sequences that successfully ran on all methods were used. In addition, we attempted to emulate the evaluation strategy of the CAID competitions. In particular, when reporting F1 and MCC, we use the binary labels reported by CAID whenever available for a method since CAID identifies the threshold that maximizes F1 score for a particular method (***Necci et al., 2021***). In the case of methods where a binary label was not provided by CAID and for DR-BERT, we identify the threshhold that maximizes F1 score ourselves. The threshhold for protein binding is calculated independently from the threshold for disordered region prediction (including disorder, disorder-PDB and disorder-noX). However, the same variant of DR-BERT was used for both disordered region and protein binding prediction.

## Acknowledgments

This work utilizes resources supported by the National Science Foundation’s Major Research Instrumentation program, grant #1725729, as well as the University of Illinois at Urbana-Champaign (***Kindratenko et al., 2020***). Part of this work was performed under the auspices of the U.S. Department of Energy by Argonne National Laboratory under Contract DE-AC02-06-CH11357. JMF and SL have been supported by the James Scholar Honors Program and the Illinois Scholars Undergraduate Research Program. We thank Mark Hopkins, Anna Ritz, Ashley Blystone and Desiree Odgers for insightful discussions.

## Author Contributions

All authors designed the study. S.M. supervised the study; A.N., J.M.F. and S.L. performed simulations and calculations. All authors discussed and wrote the paper.

## References

Alderson TR, Pritišanac I, Moses AM, Forman-Kay JD. Systematic identification of conditionally folded intrinsically disordered regions by AlphaFold2. bioRxiv. 2022; https://www.biorxiv.org/content/early/2022/02/18/2022.02.18.481080, doi: 10.1101/2022.02.18.481080.

Asgari E, Mofrad MR. Continuous distributed representation of biological sequences for deep proteomics and genomics. PloS one. 2015; 10(11):e0141287.

Bhattacharya N, Thomas N, Rao R, Dauparas J, Koo PK, Baker D, Song YS, Ovchinnikov S. Single Layers of Attention Suffice to Predict Protein Contacts. bioRxiv. 2020; https://www.biorxiv.org/content/early/2020/12/22/2020.12.21.423882.1, doi: 10.1101/2020.12.21.423882.

Campen A, Williams RM, Brown CJ, Meng J, Uversky VN, Dunker AK. TOP-IDP-scale: a new amino acid scale measuring propensity for intrinsic disorder. Protein and peptide letters. 2008; 15(9):956–963.

Chicco D, Jurman G. The advantages of the Matthews correlation coefficient (MCC) over F1 score and accuracy in binary classification evaluation. BMC genomics. 2020; 21:1–13.

Davis J, Goadrich M. The relationship between Precision-Recall and ROC curves. In: Proceedings of the 23rd international conference on Machine learning; 2006. p. 233–240.

Deng X, Eickholt J, Cheng J. PreDisorder: ab initio sequence-based prediction of protein disordered regions. BMC bioinformatics. 2009; 10:1–6.

Devlin J, Chang M, Lee K, Toutanova K. BERT: Pre-training of Deep Bidirectional Transformers for Language Understanding. CoRR. 2018; abs/1810.04805. http://arxiv.org/abs/1810.04805.

Fischer E. Einfluss der Configuration auf die Wirkung der Enzyme. Berichte der deutschen chemischen Gesellschaft. 1894; 27(3):2985–2993.

Fu L, Niu B, Zhu Z, Wu S, Li W. CD-HIT: accelerated for clustering the next-generation sequencing data. Bioin-formatics. 2012; 28(23):3150–3152.

Hanson J, Paliwal KK, Litfin T, Zhou Y. SPOT-Disorder2: improved protein intrinsic disorder prediction by en-sembled deep learning. Genomics, proteomics & bioinformatics. 2019; 17(6):645–656.

Hie BL, Yang KK, Kim PS. Evolutionary velocity with protein language models predicts evolutionary dynamics of diverse proteins. Cell Systems. 2022; 13(4):274–285.e6. https://www.sciencedirect.com/science/article/pii/S2405471222000382, doi: https://doi.org/10.1016/j.cels.2022.01.003.

Hu G, Katuwawala A, Wang K, Wu Z, Ghadermarzi S, Gao J, Kurgan L. flDPnn: Accurate intrinsic disorder predic-tion with putative propensities of disorder functions. Nature communications. 2021; 12(1):4438.

Ilzhöfer D, Heinzinger M, Rost B. SETH predicts nuances of residue disorder from protein embeddings. Fron-tiers in Bioinformatics. 2022; 2.

Ishiguro A, Nogi Y, Hisatake K, Muramatsu M, Ishihama A. The Rpb6 subunit of fission yeast RNA polymerase II is a contact target of the transcription elongation factor TFIIS. Molecular and Cellular Biology. 2000; 20(4):1263– 1270.

Kindratenko V, Mu D, Zhan Y, Maloney J, Hashemi SH, Rabe B, Xu K, Campbell R, Peng J, Gropp W. In: HAL: Computer System for Scalable Deep Learning New York, NY, USA: Association for Computing Machinery; 2020. p. 41–48. https://doi.org/10.1145/3311790.3396649.

Liu Y, Ott M, Goyal N, Du J, Joshi M, Chen D, Levy O, Lewis M, Zettlemoyer L, Stoyanov V. RoBERTa: A Robustly Optimized BERT Pretraining Approach. CoRR. 2019; abs/1907.11692. http://arxiv.org/abs/1907.11692.

Lupo U, Sgarbossa D, Bitbol AF. Protein language models trained on multiple sequence alignments learn phy-logenetic relationships. Nature Communications. 2022; 13(1):6298.

Van der Maaten L, Hinton G. Visualizing data using t-SNE. Journal of machine learning research. 2008; 9(11).

Meier J, Rao R, Verkuil R, Liu J, Sercu T, Rives A. Language models enable zero-shot prediction of the effects of mutations on protein function. In: Ranzato M, Beygelzimer A, Dauphin Y, Liang PS, Vaughan JW, editors. Advances in Neural Information Processing Systems, vol. 34 Curran Associates, Inc.; 2021. p. 29287–29303. https://proceedings.neurips.cc/paper_files/paper/2021/file/f51338d736f95dd42427296047067694-Paper.pdf.

Mészáros B, Erdős G, Dosztányi Z. IUPred2A: context-dependent prediction of protein disorder as a function of redox state and protein binding. Nucleic acids research. 2018; 46(W1):W329–W337.

Mirabello C, Wallner B. rawMSA: End-to-end deep learning using raw multiple sequence alignments. PloS one. 2019; 14(8):e0220182.

Mészáros B, Simon I, Dosztányi Z. Prediction of Protein Binding Regions in Disordered Proteins. PLOS Computational Biology. 2009 05; 5(5):1–18. https://doi.org/10.1371/journal.pcbi.1000376, doi: 10.1371/jour-nal.pcbi.1000376.

Nambiar A, Heflin M, Liu S, Maslov S, Hopkins M, Ritz A. Transforming the language of life: transformer neural networks for protein prediction tasks. In: Proceedings of the 11th ACM international conference on bioinformat-ics, computational biology and health informatics; 2020. p. 1–8.

Necci M, Piovesan D, Tosatto SC. Critical assessment of protein intrinsic disorder prediction. Nature methods. 2021; 18(5):472–481.

Okuda M, Suwa T, Suzuki H, Yamaguchi Y, Nishimura Y. Three human RNA polymerases interact with TFIIH via a common RPB6 subunit. Nucleic Acids Research. 2021 07; 50(1):1–16. https://doi.org/10.1093/nar/gkab612, doi: 10.1093/nar/gkab612.

Orlando G, Raimondi D, Codice F, Tabaro F, Vranken W. Prediction of disordered regions in proteins with recurrent neural networks and protein dynamics. Journal of Molecular Biology. 2022; 434(12):167579.

Piovesan D, Tabaro F, Micetic I, Necci M, Quaglia F, Oldfield CJ, Aspromonte MC, Davey NE, Davidovic R, Dosztányi Z, Elofsson A, Gasparini A, Hatos A, Kajava AV, Kalmar L, Leonardi E, Lazar T, Macedo-Ribeiro S, Macossay-Castillo M, Meszaros A, et al. DisProt 7.0: a major update of the database of disordered proteins. Nucleic Acids Research. 2016 11; 45(D1):D219–D227. https://doi.org/10.1093/nar/gkw1056, doi: 10.1093/nar/gkw1056.

Quaglia F, Mészáros B, Salladini E, Hatos A, Pancsa R, Chemes LB, Pajkos M, Lazar T, Peña-Díaz S, Santos J, Ács V, Farahi N, Fichó E, Aspromonte M, Bassot C, Chasapi A, Davey N, Davidovic R, Dobson L, Elofsson A, et al. DisProt in 2022: improved quality and accessibility of protein intrinsic disorder annotation. Nucleic Acids Research. 2021 11; 50(D1):D480–D487. https://doi.org/10.1093/nar/gkab1082, doi: 10.1093/nar/gkab1082.

Redl I, Fisicaro C, Dutton O, Hoffmann F, Henderson L, Owens BMJ, Heberling M, Paci E, Tamiola K. ADOPT: intrinsic protein disorder prediction through deep bidirectional transformers. bioRxiv. 2023; https://www.biorxiv.org/content/early/2023/01/11/2022.05.25.493416, doi: 10.1101/2022.05.25.493416.

Rives A, Meier J, Sercu T, Goyal S, Lin Z, Liu J, Guo D, Ott M, Zitnick CL, Ma J, Fergus R. Biological structure and function emerge from scaling unsupervised learning to 250 million protein sequences. Proceedings of the National Academy of Sciences. 2021; 118(15):e2016239118. https://www.pnas.org/doi/abs/10.1073/pnas. 2016239118, doi: 10.1073/pnas.2016239118.

Singh J, Paliwal K, Litfin T, Singh J, Zhou Y. Reaching alignment-profile-based accuracy in predicting protein secondary and tertiary structural properties without alignment. Scientific Reports. 2022; 12(1):7607.

Srivastava N, Hinton G, Krizhevsky A, Sutskever I, Salakhutdinov R. Dropout: A Simple Way to Prevent Neural Networks from Overfitting. Journal of Machine Learning Research. 2014; 15(56):1929–1958. http://jmlr.org/papers/v15/srivastava14a.html.

Stärk H, Dallago C, Heinzinger M, Rost B. Light attention predicts protein location from the lan-guage of life. Bioinformatics Advances. 2021 11; 1(1). https://doi.org/10.1093/bioadv/vbab035, doi: 10.1093/bioadv/vbab035, vbab035.

Suzek BE, Wang Y, Huang H, McGarvey PB, Wu CH, the UniProt Consortium. UniRef clusters: a comprehensive and scalable alternative for improving sequence similarity searches. Bioinformatics. 2014 11; 31(6):926–932. https://doi.org/10.1093/bioinformatics/btu739, doi: 10.1093/bioinformatics/btu739.

Tang YJ, Pang YH, Liu B. IDP-Seq2Seq: identification of intrinsically disordered regions based on sequence to sequence learning. Bioinformatics. 2020; 36(21):5177–5186.

Tang YJ, Pang YH, Liu B. DeepIDP-2L: protein intrinsically disordered region prediction by combining convolu-tional attention network and hierarchical attention network. Bioinformatics. 2022; 38(5):1252–1260.

Uversky VN. Natively unfolded proteins: A point where biology waits for physics. Protein Science. 2002; 11(4):739–756. https://onlinelibrary.wiley.com/doi/abs/10.1110/ps.4210102, doi: https://doi.org/10.1110/ps.4210102.

Van Der Lee R, Buljan M, Lang B, Weatheritt RJ, Daughdrill GW, Dunker AK, Fuxreiter M, Gough J, Gsponer J, Jones DT, et al. Classification of intrinsically disordered regions and proteins. Chemical reviews. 2014; 114(13):6589–6631.

Vaswani A, Shazeer N, Parmar N, Uszkoreit J, Jones L, Gomez AN, Kaiser Lu, Polosukhin I. Attention is All you Need. In: Guyon I, Luxburg UV, Bengio S, Wallach H, Fergus R, Vishwanathan S, Garnett R, editors. Advances in Neural Information Processing Systems, vol. 30 Curran Associates, Inc.; 2017. https://proceedings.neurips.cc/paper/2017/file/3f5ee243547dee91fbd053c1c4a845aa-Paper.pdf.

Walsh I, Martin AJ, Di Domenico T, Tosatto SC. ESpritz: accurate and fast prediction of protein disorder. Bioin-formatics. 2012; 28(4):503–509.

Wang S, Ma J, Xu J. AUCpreD: proteome-level protein disorder prediction by AUC-maximized deep convolutional neural fields. Bioinformatics. 2016; 32(17):i672–i679.

Wright PE, Dyson HJ. Intrinsically disordered proteins in cellular signalling and regulation. Nature reviews Molecular cell biology. 2015; 16(1):18–29.

Wright PE, Dyson HJ. Intrinsically unstructured proteins: re-assessing the protein structure-function paradigm. Journal of Molecular Biology. 1999; 293(2):321–331. https://www.sciencedirect.com/science/article/pii/S0022283699931108, doi: https://doi.org/10.1006/jmbi.1999.3110.

Zhao B, Kurgan L. Deep learning in prediction of intrinsic disorder in proteins. Computational and Structural Biotechnology Journal. 2022; 20:1286–1294. https://www.sciencedirect.com/science/article/pii/S2001037022000782, doi: https://doi.org/10.1016/j.csbj.2022.03.003.

