## Supplementary Figures for "DR-BERT: A Protein Language Model to Annotate Disordered Regions"

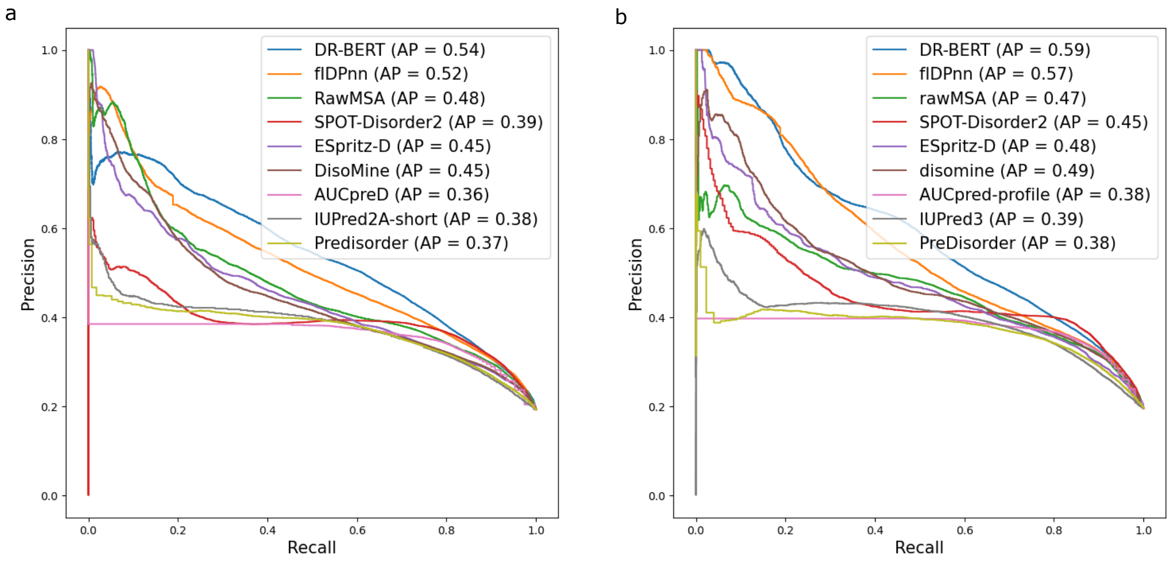


SUPP.FIG. 1. **The Precision-Recall plots of DR-BERT and the top performing models from (a) CAID 1 and (b) CAID 2.** The area under each curve (AP) is presented in the legend.

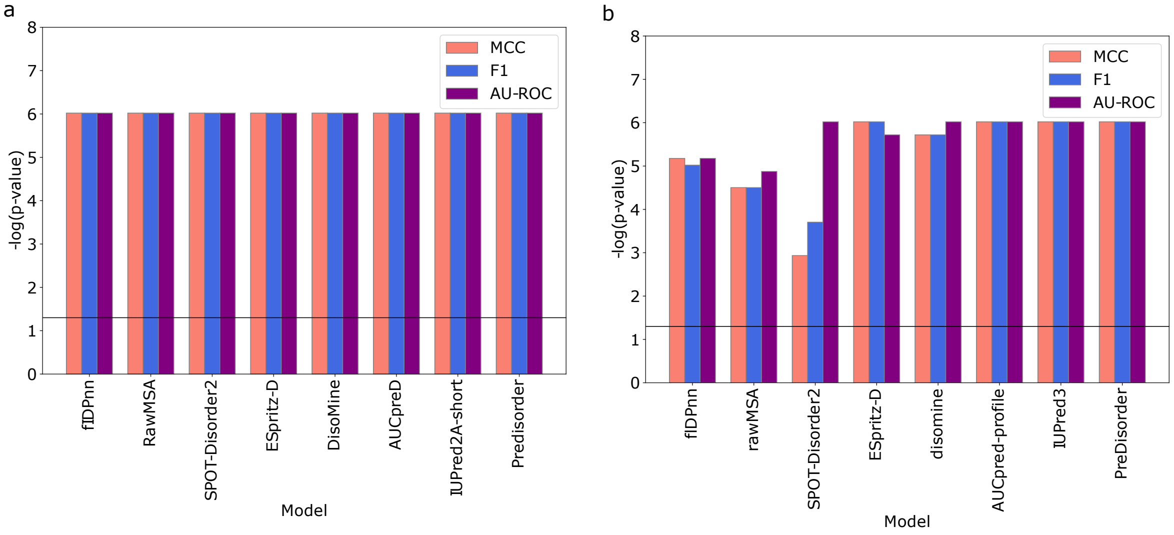


SUPP.FIG. 2. **The results from resampling the test set for (a) CAID 1 and (b) CAID 2.** These plots show the $\log_{10}$ of p-value of the Wilcoxon tests comparing DR-BERT to the other methods. The horizontal line indicates the 0.05 p-value threshold, above which DR-BERT significantly outperforms the other methods.

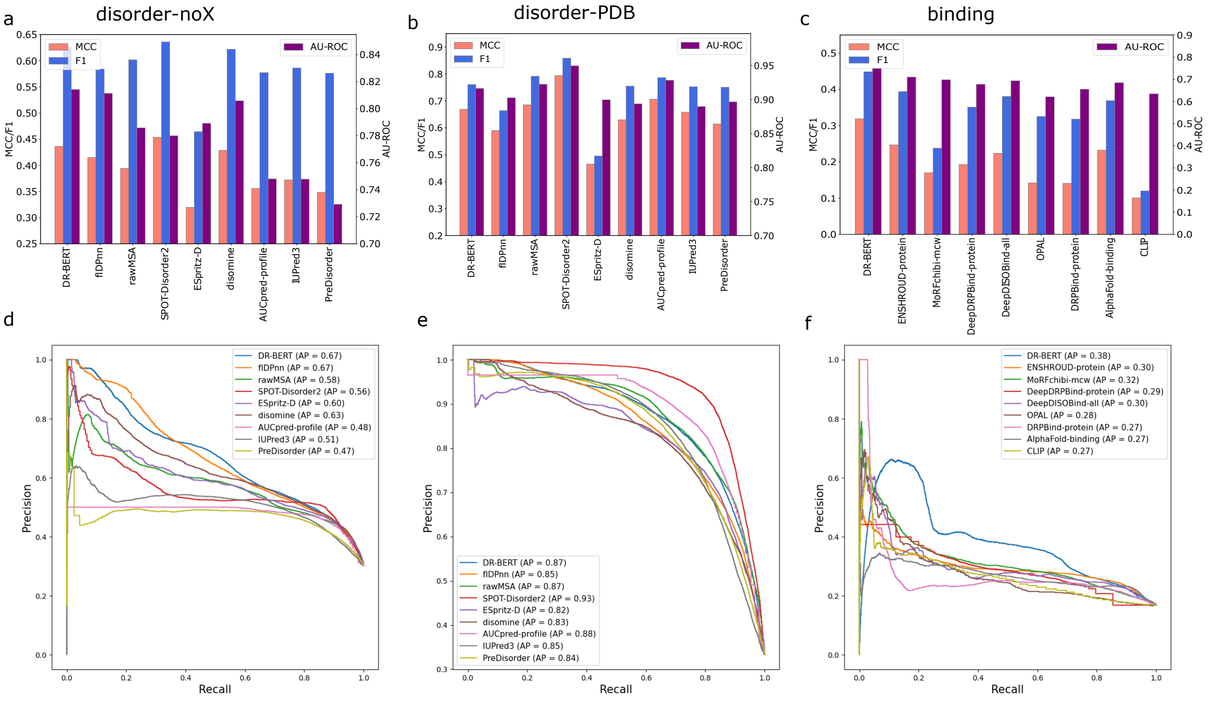


SUPP.FIG. 3. **The results on the additional test sets from CAID 2. (a, b and c)** Show the AU-ROC, F1 score and MCC on the disorder-noX, disorder-PDB and protein binding test sets. **(d,e, f)** Show precision recall plots that correspond to the same tasks.

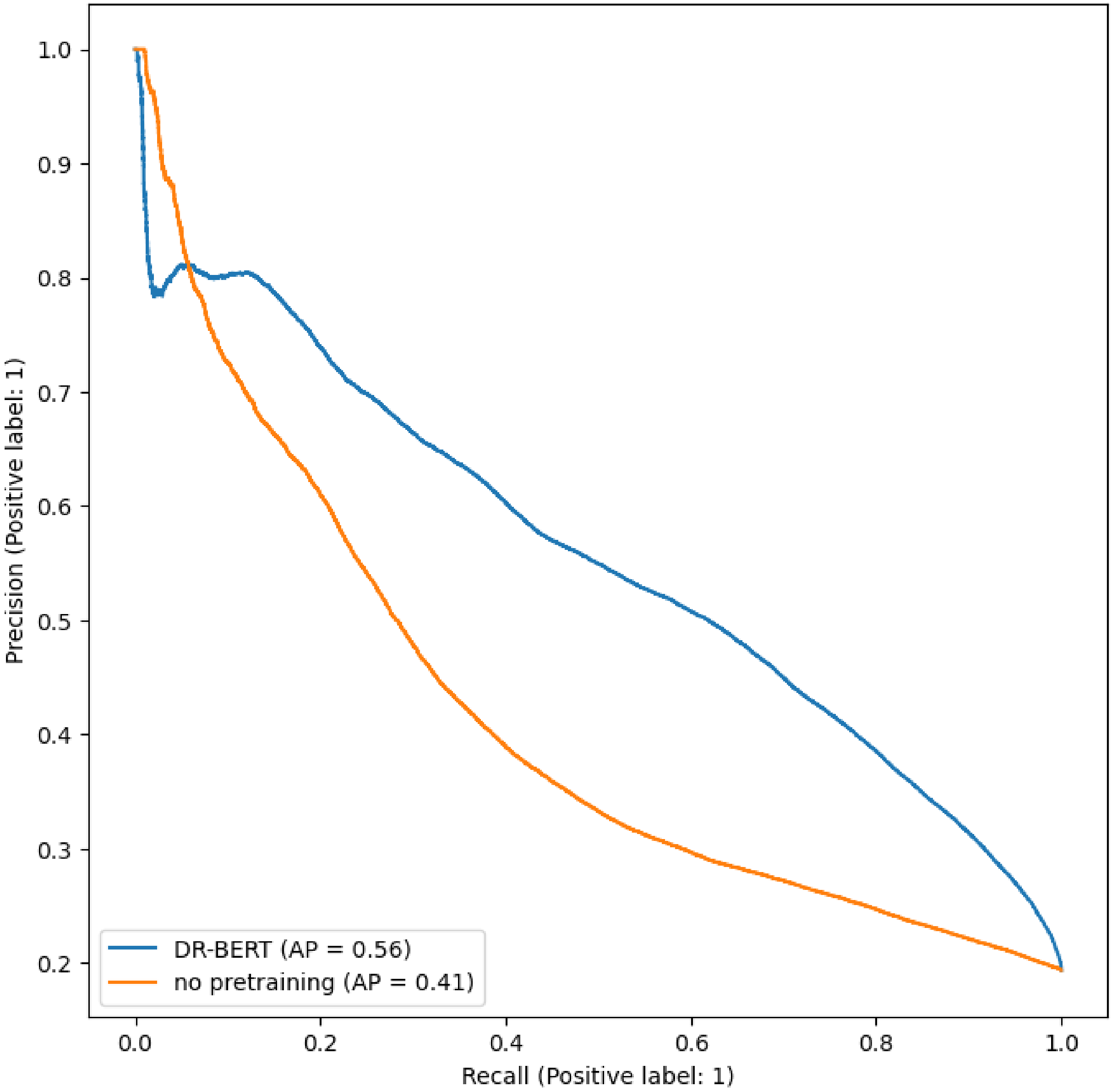


SUPP.FIG. 4. **Comparing DR-BERT to a non-pretrained version.** The Precision-Recall plots of DR-BERT and the non-pretrained model. The area under each curve (AP) is presented in the legend

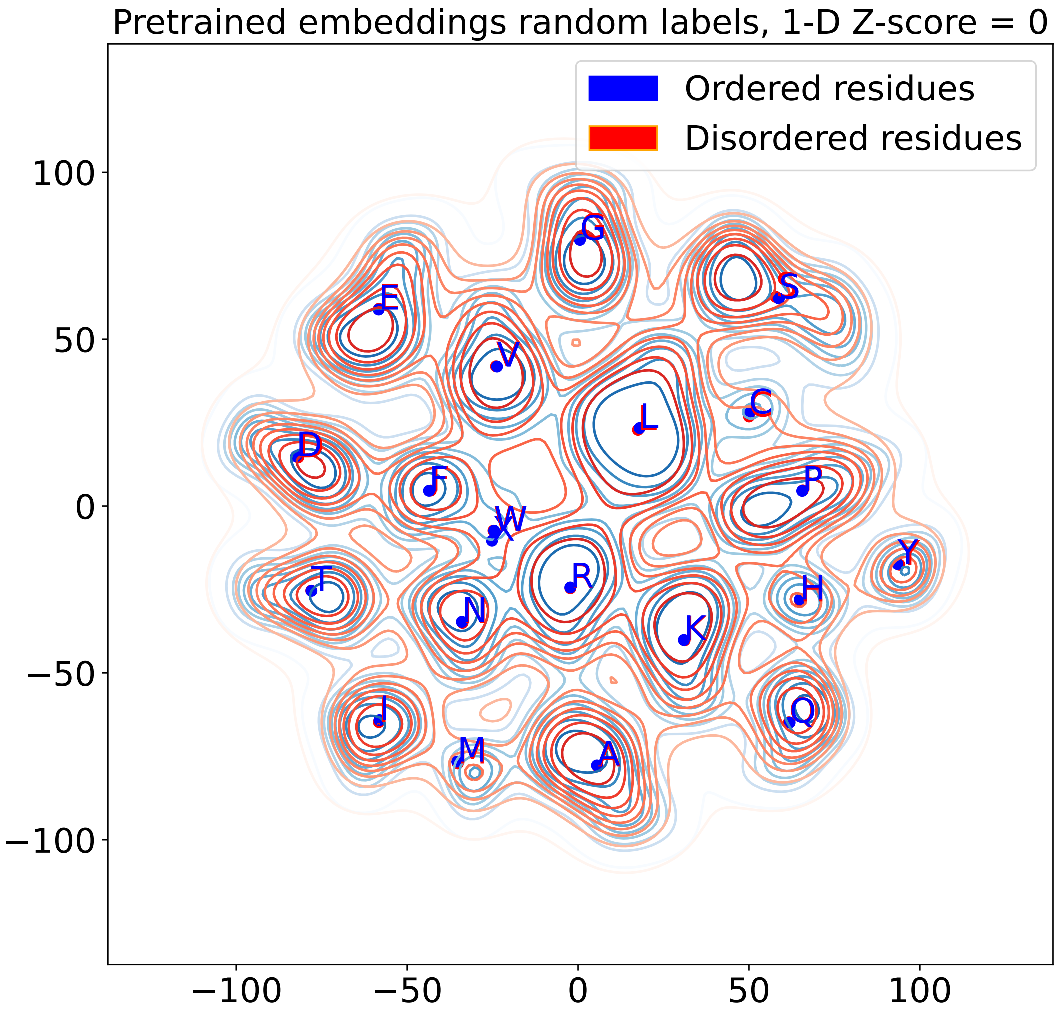


SUPP.FIG. 5. **A null model for the pretrained embeddings** A t-SNE projection of the pretrained embeddings of residues in the test set for CAID 1. The disorder/order residue labels have been shuffled here to create a null model.

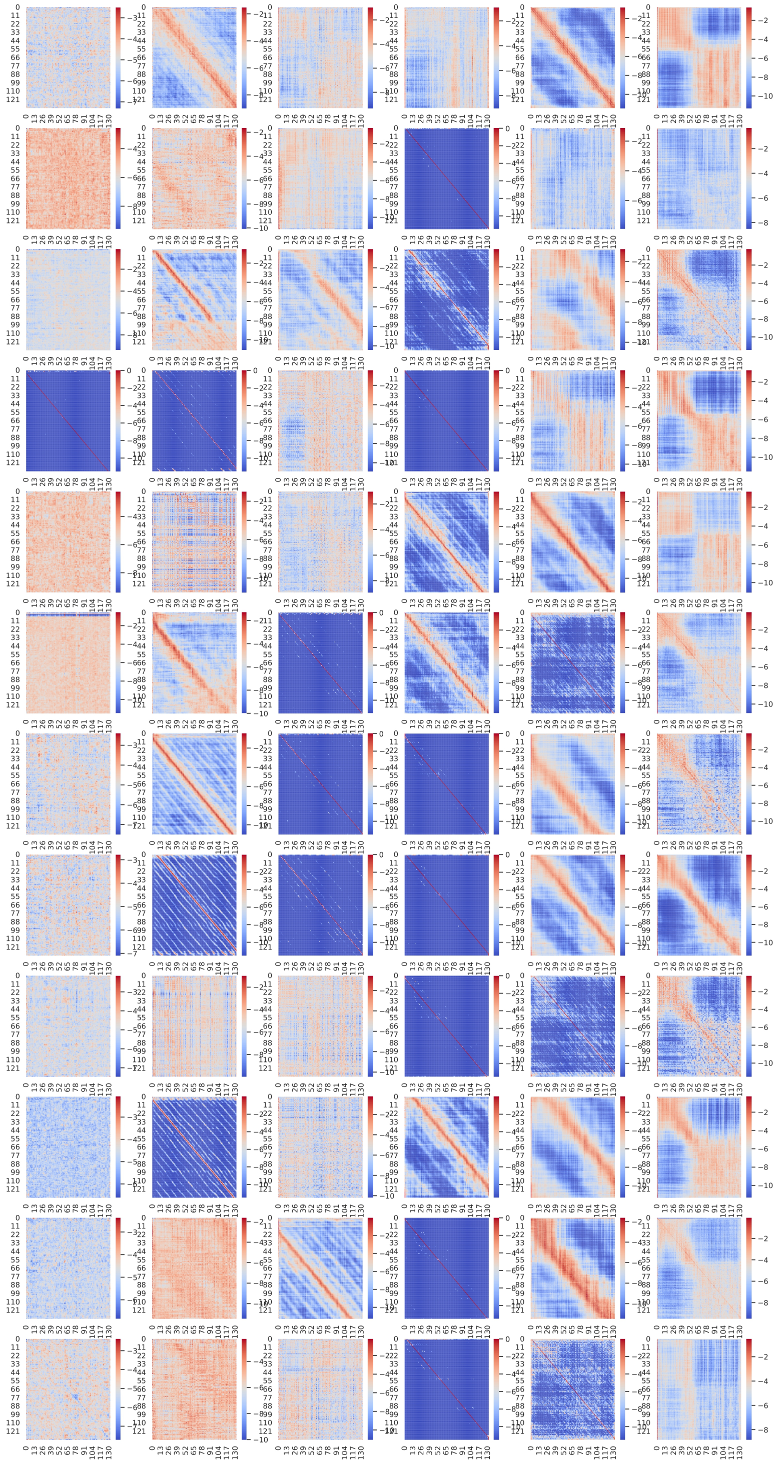

SUPP.FIG. 6. **Vizualizing the attention heads of DR-BERT as it processes the RPB6 protein.** Each of the 12 attention maps for each of the 6 encoder layers are shown here. The columns correspond to different encoder layers with the left-most column being the first layer and the right-most being the sixth layer.

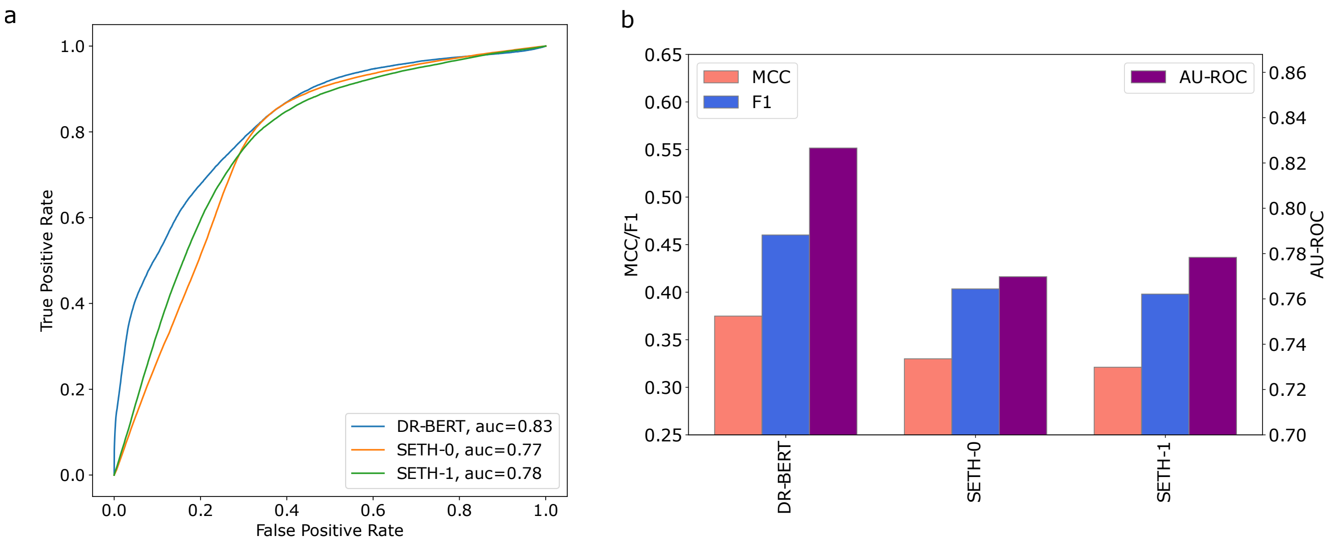


SUPP.FIG. 7. **Comparing DR-BERT to a another Transformer based model, SETH. (a)** The ROC plots of DR-BERT and two versions of SETH on CAID 2. The area under each curve (AUC) is presented in the legend. **(b)** The MCC, F1 and AU-ROC scores of DR-BERT and the two SETH variants.
